# MARK1 regulates dendritic spine morphogenesis and cognitive functions *in vivo*

**DOI:** 10.1101/2023.12.03.569757

**Authors:** Emily C. Kelly-Castro, Rebecca Shear, Ankitha H. Dindigal, Maitreyee Bhagwat, Huaye Zhang

**Author notes:** Corresponding author: Dr. Huaye Zhang, Department of Neuroscience and Cell Biology, Rutgers Robert Wood Johnson Medical School, 675 Hoes Lane West, Piscataway, NJ 08854. These authors contributed equally (Emily C. Kelly-Castro, Rebecca Shear).

## Abstract

Dendritic spines play a pivotal role in synaptic communication and are crucial for learning and memory processes. Abnormalities in spine morphology and plasticity are observed in neurodevelopmental and neuropsychiatric disorders, yet the underlying signaling mechanisms remain poorly understood. The microtubule affinity regulating kinase 1 (MARK1) has been implicated in neurodevelopmental disorders, and the *MARK1* gene shows accelerated evolution in the human lineage suggesting a role in cognition. However, the *in vivo* role of MARK1 in synaptogenesis and cognitive functions remains unknown. Here we show that forebrain-specific conditional knockout (cKO) of *Mark1* causes defects in dendritic spine morphogenesis in hippocampal CA1 pyramidal neurons with a significant reduction in spine density. In addition, we found that MARK1 cKO mice show defects in spatial learning in the Morris Water Maze and reduced anxiety-like behaviors in the Elevated Plus Maze. Furthermore, we found loss of MARK1 causes synaptic accumulation of GKAP and GluR2. Taken together, our data show a novel role for MARK1 in regulating dendritic spine morphogenesis and cognitive functions *in vivo*.

## Introduction

Dendritic spines are small actin-rich protrusions on mammalian neurons that act as the primary receivers of excitatory synaptic inputs^1-5^. Spines are dynamic, and they can change in size and shape depending on the input signaling received. The size of the spine head is usually proportional to the postsynaptic density (PSD), which is an electron-dense zone underneath the postsynaptic membrane that contains receptors, scaffolding proteins, and cytoskeletal proteins, among others^6^. The enlargement of dendritic spines and their associated PSD correlates with long-term potentiation (LTP), a long-lasting increase in the efficacy of neurotransmission^7-9^.

Conversely, spine shrinkage correlates with a decrease in neurotransmission known as long-term depression (LTD)^10-13^. This structural plasticity of dendritic spines underlies learning and memory processes^14^. Two-photon *in vivo* imaging studies have revealed that learning is associated with the formation of new spines and elimination of existing spines in a region- and circuit-specific manner^15-17^. Optogenetic shrinkage of newly formed dendritic spines after motor learning erases task-specific memories, suggesting a causal relationship between dendritic spine structural plasticity and learning^18^. Not surprisingly, abnormalities in spine morphology and plasticity are observed in a number of neurodevelopmental and neuropsychiatric disorders, including autism spectrum disorders (ASD), schizophrenia, and bipolar disorder^19-21^. Previous studies show that spine density increases in the brains of autistic patients^22,23^, and decreases in the brains of schizophrenic patients^24^. However, our understanding of the signaling mechanisms that lead to spine abnormalities and cognitive dysfunctions remains limited.

We and others previously showed that the microtubule affinity regulating kinase (MARK) family of kinases, also known as partitioning defective 1 (Par1), are involved in dendritic spine morphogenesis and structural plasticity in primary hippocampal neurons^4,25-27^. We found that knockdown of either MARK1 or MARK2 results in defects in spine morphogenesis. Moreover, elevated expression of MARK kinases also inhibits the proper formation of dendritic spines, indicating that precise spatiotemporal control of MARK family kinases is necessary^4^. We further found that MARK kinases are activated downstream of N-methyl-D-aspartate (NMDA) receptors, which are glutamatergic ionotropic receptors on dendritic spines that play key roles in synaptic plasticity^26^. Finally, we found that MARK kinases function by phosphorylating the postsynaptic scaffolding protein PSD-95 on Ser561^4^. Phosphorylation by MARK leads to a conformational switch within PSD-95 that decreases its interaction with its binding partners such as GKAP and increases PSD-95 dynamics. In addition, a non-phosphorylatable mutant of PSD-95 Ser561 inhibits bidirectional dendritic spine structural plasticity, indicating that MARK-mediated phosphorylation of PSD-95 is critical for synaptic plasticity^27^.

Interestingly, multiple lines of evidence point to an important role of MARK family kinases in higher-order cognitive functions. *MARK1* shows highly accelerated evolution in the human lineage, with a Ka/Ks ratio (non-synonymous vs. synonymous substitution rates) of 2.11 when comparing humans with chimpanzees^28^. Notably, fewer than 1% of the genes have a Ka/Ks>2, as the vast majority of genes are under purifying selection, with an average Ka/Ks of 0.23 between humans and chimpanzees^29^. This adaptive evolution of *MARK1*, combined with its high expression in the brain, suggests a possible role of this gene in human-specific cognitive functions^28^. Moreover, multiple SNPs have been found in *MARK1* that are associated with autism. One of the SNPs has been shown to modulate *MARK1* transcription, and *MARK1* mRNA was overexpressed in the prefrontal cortex of autistic patients^28^. Furthermore, MARK1 was also found to be associated with bipolar disorder^30^, which points to a general involvement of this kinase in psychiatric disorders. However, the role for MARK1 in cognitive functions *in vivo* is completely unknown.

To examine the role of MARK1 in dendritic spine morphogenesis and cognitive functions *in vivo*, we established floxed *Mark1* alleles and crossed these mice with *CaMKIIα-Cre* mice to knock out *Mark1* from postnatal forebrain pyramidal neurons (*Mark1*^*f/f*^*:CaMKIIα-Cre*). Here we show that forebrain specific conditional knockout (cKO) of MARK1 causes defects in dendritic spine morphogenesis in hippocampal CA1 pyramidal neurons with a significant reduction in spine density. In addition, we found that MARK1 cKO mice show defects in spatial learning in the Morris Water Maze and reduced anxiety-like behaviors in the Elevated Plus Maze.

Furthermore, we found loss of MARK1 causes synaptic accumulation of GKAP and GluR2. Taken together, our data show a novel role for MARK1 in regulating dendritic spine morphogenesis and cognitive functions *in vivo*.

## Materials and Methods

### Animals

All animals used in this study were carried out in accordance with Rutgers Institutional Animal Care and Use Committee (IACUC) protocols. Floxed *Mark1* alleles were generated by Biocytogen through inserting LoxP sites flanking exon 4 of *Mark1* using CRISPR/Cas9. Mice were backcrossed with C57/Bl6 mice for at least five generations before experiments were initiated. B6. Cg-Tg (CaMKIIa-Cre) T29-1Stl/J mice were purchased from Jackson Laboratories (Strain #005359). Forebrain-specific conditional knockout (cKO) of *Mark1* was achieved by crossing floxed *Mark1* mice with B6. Cg-Tg (CaMKIIa-Cre) T29-1Stl/J mice. This will knock out *Mark1* in forebrain pyramidal neurons starting around 3 weeks postnatally^31^. Mice were housed with free access to food and water in a temperature and humidity-regulated room with a 12hr light/dark cycle.

### Genotyping

The genotypes of the MARK1 cKO mice were determined with PCR using the AccuStartII genotyping kit (Quantabio, Beverly, MA). The following oligonucleotides were used: primer 1 MARK1-5’loxP-F (5’-CTCATCCTCTTGCCCAGCCTTATG-3’), and primer 2 MARK1-5’loxP-R (5’-AGAAGAAAAGCGGGAGGAAATGGCA-3’). Primers 1 and 2 amplified a 242-bp fragment specific to the wild-type allele and a 341-bp fragment specific to the cKO allele. The genotypes of the CaMKIIa-Cre mice were determined with PCR using the following oligonucleotides: primer 1 Cre GT-F (5’-GAACCTGATGGACATGTTCAGG-3’) and primer 2 Cre GT-R (5’-AGTGCGTTCGAACGCTAGAGCCTGT-3’). Primers 1 and 2 amplified a 300-bp fragment specific to the *cre* allele.

### Golgi Staining

Brains were taken from six-week-old mice and processed using the FD Neuro Technologies Rapid Golgi kit (#PK401, FD NeuroTechnologies, MD). Briefly, animal brains were quickly removed, rinsed 2x in Milli-Q water, and rapidly immersed into the impregnation solution. The solution was replaced with fresh impregnation solution on the next day, and the tissue was stored at room temperature for three weeks in the dark. After three weeks, the brain tissue was transferred into solution C and stored at room temperature for 72 hours in the dark. After 72 hours of incubation in solution C, brains were quickly frozen using pre-cooled isopentane and stored at -80°C until sectioning. Brains were sectioned to 150μm and mounted on 2% gelatin slides. After sectioning the brains, the samples were fully processed within three days of sectioning following the manufacturer’s protocol. Z-stack images (1μm step) were taken of the CA1 apical dendrites from each group.

### Microscopy and image analysis

A Leica bright field microscope was used to acquire the z-stack images. Apical dendrites of pyramidal neurons from the CA1 region were imaged using a 63X oil lens. Spine density, length, and width were measured using FIJI and RECONSTRUCT. Spine density was calculated as the number of dendritic protrusions per 100μm. The length of the spines was measured from the starting point of the neck of the spine to the tip of the head of the spine.

Spine width was measured as the widest point of the spine head perpendicular to the length measurement. Spines were then classified into shape categories based on the length and width measurements: mushroom, ***H/N*** > 1.2, 0.6 μm ***< L*** < 1 μm; stubby, ***H/N*** ≤*1*.*2*, ***W*** ≤ 0.6 μm; long thin, ***H/N*** > 1.2, 1 μm **≤ *L*** < 2; filipodia **L** ≥ 2 μm. Spines in each morphological category were quantified by spine count per 10μm dendrite.

### Synaptosome preparation and Western Blot

Mice were euthanized by cervical dislocation, and brains were quickly removed, immediately washed with 1x PBS to remove excess blood, and transferred into a glass petri dish with cold 1x PBS to dissect the cortex and the hippocampus. Each piece of tissue was transferred into a 1.5ml centrifuge tube and rapidly snap-frozen in liquid nitrogen. Samples were then stored at - 80°C until further processing. An electric hand-held homogenizer was used to homogenize the tissue in lysis buffer 1 containing 10mM Hepes, pH 7.4, 2mM EDTA, protease inhibitor 1:1000, and phosphatase inhibitor 1:100. Lysates were cleared by centrifugation at 1,000g for 10 minutes at 4°C. From the resulting supernatant, fractions of 20μl were saved and labeled as homogenate for further analysis. The remaining supernatant was transferred into a new tube and further cleared by centrifugation at 12,000g for 20 minutes at 4°C. After this step, the supernatant was transferred into a new tube, labeled as S1, the cytosolic/membrane fraction, and saved for further analysis. The remaining pellet is the crude synaptosome fraction resuspended using buffer 2 containing 50mM HEPES, 2mM EDTA, 2mM EGTA, 1% Triton X-100, protease inhibitor 1:1000, and phosphatase inhibitor 1:100, and saved at -80°C for further analysis. The primary antibodies used were rabbit anti-MARK1 (1:1000, Proteintech, Chicago, IL), rabbit-anti-MARK2 (1:1000, Cell Signaling Tech, Boston, MA), mouse anti-MARK3 (C-TAK) (1:1000, EMD Millipore, Temecula, CA), rabbit anti-MARK4 (1:1000, Santa Cruz Biotechnology, Santa Cruz, CA), mouse anti-GluR1(1:1000, Neuromab, Davis, CA), mouse anti-GluR2(1:1000, Neuromab, Davis, CA), mouse anti-NMDAR1(1:1000, Neuromab, Davis, CA), rabbit anti-NMDAR2B (1:1000, Bioss USA, Woburn, MA), rabbit anti-mGluR5 (1:1000, Upstate Bio, Temecula, CA), mouse anti-PSD-95 (1:1000, 6G6, Santa Cruz Biotechnology, Santa Cruz, CA), mouse anti-Pan-SAPAP (1:1000, Neuromab, Davis, CA), mouse anti-SHANK3 (1:1000, Neuromab, Davis, CA), and mouse anti-GAPDH. Horseradish peroxidase-conjugated goat anti-rabbit and goat anti-mouse secondary antibodies (Jackson ImmunoResearch Laboratories, West Grove, PA) were used. Proteins were visualized by enhanced chemiluminescence using ECL Western blotting substrate and imaged on an Azure c600 (Azure Biosystems) imaging station and software (Version 1.4.0.0309; AI600-1090).

### Mouse Behavior

Morris Water Maze (MWM): The Morris Water Maze apparatus consisted of a circular, plastic pool 1.36 m in diameter (1.2 m diameter inner circle). The pool was filled with regular tap water made opaque by adding white non-toxic paint, and the temperature was kept at 21 ± 1°C. A tall platform was placed in one of the quadrants. This platform consisted of a small 10 cm, clear, perforated, plexiglass disc mounted onto a 38 cm tall steel rod and affixed to a heavy metal base. The top of the platform was hidden 1cm below the water. Visual cues were placed around the pool and hung on the walls surrounding the pool. MARK1 wildtype (WT, *Mark1*^*f/f*^) and cKO (*Mark1*^*f/f*^*:CaMKIIα-Cre*) male mice were tested for seven consecutive days, 3 trials for each day. For each trial, a maximum swim time of 60 seconds was imposed. On Day 1 a prop was placed on the platform to help the mouse identify its location. The mouse was allowed to swim for 60 seconds; if the time was up and the mouse did not find the platform, it was placed on the platform for 15 seconds. For Days 2-6, the prop was removed. During these days the latency to the hidden platform from the different starting points was recorded. A reduction in latency to platform indicates the mouse learned the platform location. On Day 7, the platform was taken away and the total amount of time the mouse spent swimming in the targeted quadrant was recorded. The amount of time spent in the target quadrant compared to the other quadrants represents retention of spatial memory.

Elevated Plus Maze (EPM): The Elevated Plus Maze evaluates anxiety-like behavior. It consisted of a ‘plus’ shaped apparatus about 60 cm above the floor with two long closed arms (1.43 m long and 8.6 cm wide), two short open arms (69.6 cm long and 8.6 cm wide), and a central square of 8.6 × 8.6 cm. Wild-type mice would normally get anxious when exposed to elevated open spaces, therefore they will tend to spend more time on the closed arms. Each mouse was placed on the center square and observed for a 10-minute period. The number of times entered, and the time spent on the open or closed arm was recorded.

Social Test: The sociability test measures the disposition of the mice to interact with other mice. Mice were placed in a 40.6”W x 40.6”L x 47”H cm plexiglass chamber. Two stainless steel mesh cups within the chamber, 33cm in diameter and 43cm tall, were located on opposite corners of the chamber. Each mouse was given a 10-minute habituation period to explore the chamber before the trial. After 10 minutes, one age- and sex-matched MARK1 WT mouse (Stranger 1) was placed in one of the stainless-steel mesh cups, and the experimental mouse was placed in the center of the chamber. The time spent in the target chamber with Stranger 1 or the chamber with the empty cup was recorded. WT and cKO mice were tested for 10 minutes on one day. More time spent in the targeted chamber with the stranger mouse would indicate a higher level of social behavior. The time interacting with each stainless-steel mesh cup and preference index was recorded.

The social novelty test was done 3 hours after the sociability test. All experimental mice were age and sex-matched with two MARK1 WT mice to serve as the familiar (Stranger 1) and the novel stranger (Stranger 2) mice. Each mouse was placed in each of the stainless-steel mesh cups. WT and cKO mice were tested for a total time of 10 minutes. The time interacting with each mouse was recorded. If more time were spent with the Stranger 2 mouse, this would indicate a higher level of social novelty behavior.

### Statistical analysis

For Golgi analysis of dendritic spines, samples were blinded for both image collection and data analysis. Behavioral and Western blot data were analyzed in GraphPad Prism using either an unpaired t-test or ANOVA with repeated measure followed by post-hoc Tukey’s test. For dendritic spine data, a mixed model analysis with simple covariance was performed using SAS to account for data nesting within animals as previously described^32^. Data are shown as Mean ± SEM with individual data points in bar graphs, or as violin plots with thick lines as the median and dotted lines as quartiles. Statistical significance was set at p<0.05.

## Results

### MARK1 is decreased in *MARK1* conditional knockout mouse model (cKO)

Our previous studies show MARK1 plays an essential role in dendritic spine morphogenesis in rat primary hippocampal neurons^4,26,27^. To investigate the role of MARK1 in dendritic spine morphogenesis, plasticity, and cognition *in vivo*, we first established floxed alleles of *Mark1* by inserting LoxP sites flanking exon 4 of *Mark1*. Forebrain-specific conditional knockout of *MARK1* was achieved by crossing *CaMKIIa-Cre* mice with *Mark1*^*fl/fl*^ mice (**Figure 1A**). The *CaMKIIa* promoter drives the expression of Cre recombinase protein in cortical and hippocampal pyramidal neurons of the mice during the third postnatal week, which allows us to avoid potential defects in processes such as embryonic neurogenesis and neuronal migration. To confirm that MARK1 protein was knocked out, Western blot analysis was done to compare protein levels in the cortex and hippocampal tissue between *Mark1*^*fl/fl*^*:CaMKIIa-Cre* mice and littermate controls (**Figure 1B**). We observed more than an 80% decrease in the amount of MARK1 protein in both cortical and hippocampal tissue from the mutant mice as compared with their WT controls (**Figure 1D**). Since the *CaMKIIa* promoter leads to the expression of Cre only in pyramidal neurons, the remaining MARK1 protein likely originated from inhibitory neurons or glial cells. No significant changes in body weight or brain weight were observed in the MARK1 cKO mice, indicating that the gross morphological development is not affected in these mice compared with WT (**Supplementary Figure 1A and B**). In addition, we observed no significant changes in the expression of MARK2 and MARK3 (**Supplementary Figure 1C**). MARK4 is expressed at low levels in the brain and therefore cannot be reliably detected by Western blot^33^. This suggests that there is no or minimal compensation from other MARK family members.

**Figure 1:**
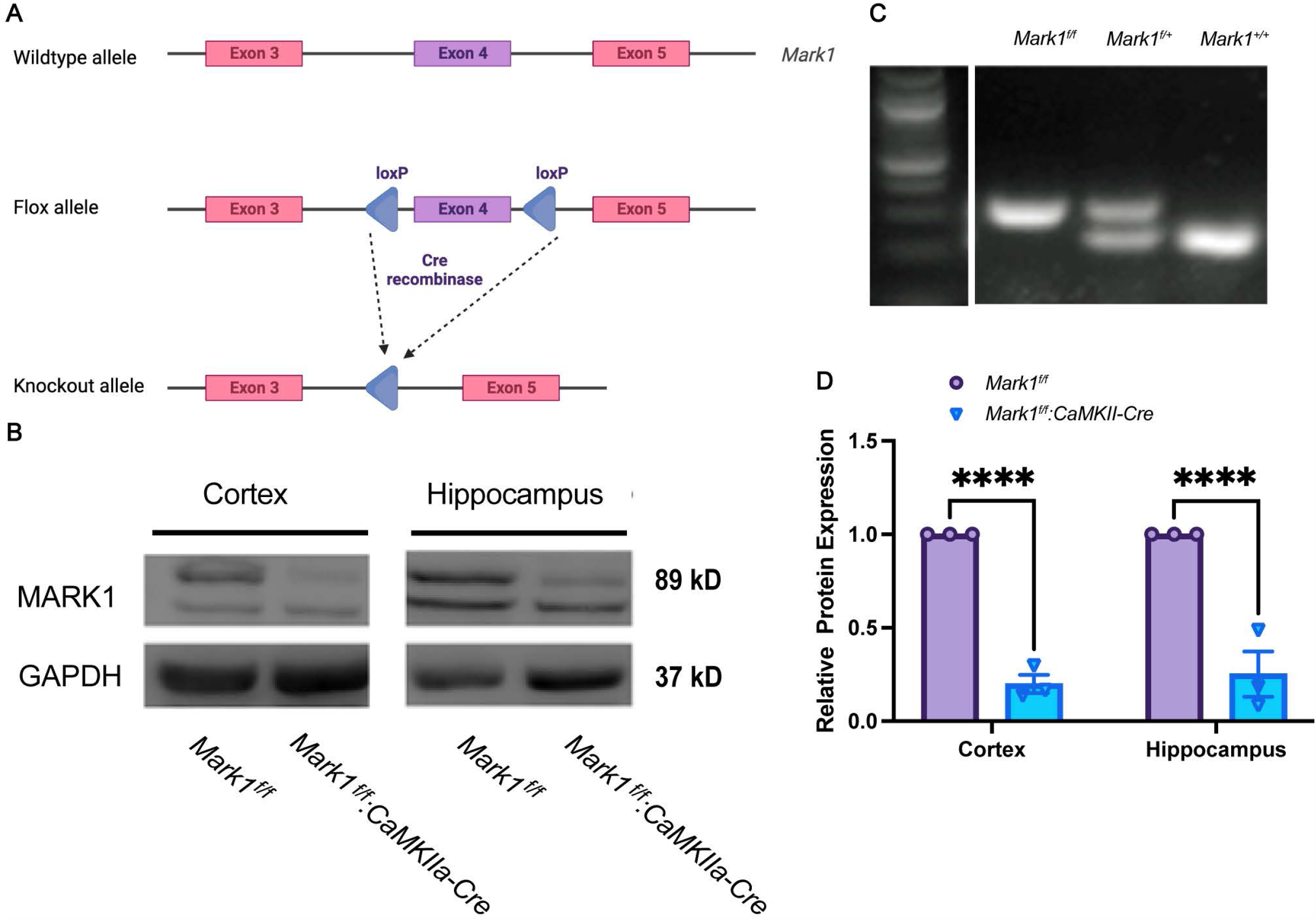
MARK1 conditional knockout mouse model. A. Schematic of *Mark1* allele where LoxP sites were inserted using CRISPR-Cas9 technology. B. Western blot image representation of MARK1 protein (∼89kDa) expression in cortical and hippocampal tissue. GAPDH (∼37kDa) was used as the loading control. C. PCR electrophoresis gel of amplified LoxP MARK1 regions. Tail samples using primers 5’F/R Lox-p MARK1 revealed bands at 341bp (*Mark1*^*f/f*^), 242bp and 341bp (*Mark1*^*f/+*^), and 242bp (*Mark1*^*+/+*^). Cre recombinase protein expression is restricted to the cortex and hippocampus. D. Graph of mean relative intensity of MARK1 protein expression. Data are shown as mean ± SEM, n=4 for *Mark1*^*f/f*^, n= 4 for *Mark1*^*f/f*^*:CaMKIIa-Cre*. Cortex and hippocampus: p<0.0001.

### Loss of MARK1 impairs dendritic spine morphogenesis in hippocampal CA1 pyramidal neurons

We next used rapid Golgi staining to analyze dendritic spines from the apical dendrites in the stratum radiatum (SR) layer of the hippocampal CA1 region in 5-week-old WT and MARK1 cKO mice. Results show that MARK1 cKO mice have a significantly reduced dendritic spine density as compared with littermate WT mice (p=0.0172) (**Figure 2A and B**). We then analyzed dendritic spine morphology by measuring spine length and spine width. We found that MARK1 cKO mice do not show a significant difference in spine length or spine width (**Figure 2C, D, and E**). Finally, we categorized dendritic spines into thin, stubby, mushroom-shaped, and branched using the morphological measurements. We found that MARK1 cKO mice show a trend towards a higher proportion of stubby shaped dendritic spines (**Figure 2F**). Together, these data show there is a reduction in the density of dendritic spines in MARK1 cKO hippocampus with a trend towards an increase in the proportion of stubby shaped spines.

**Figure 2:**
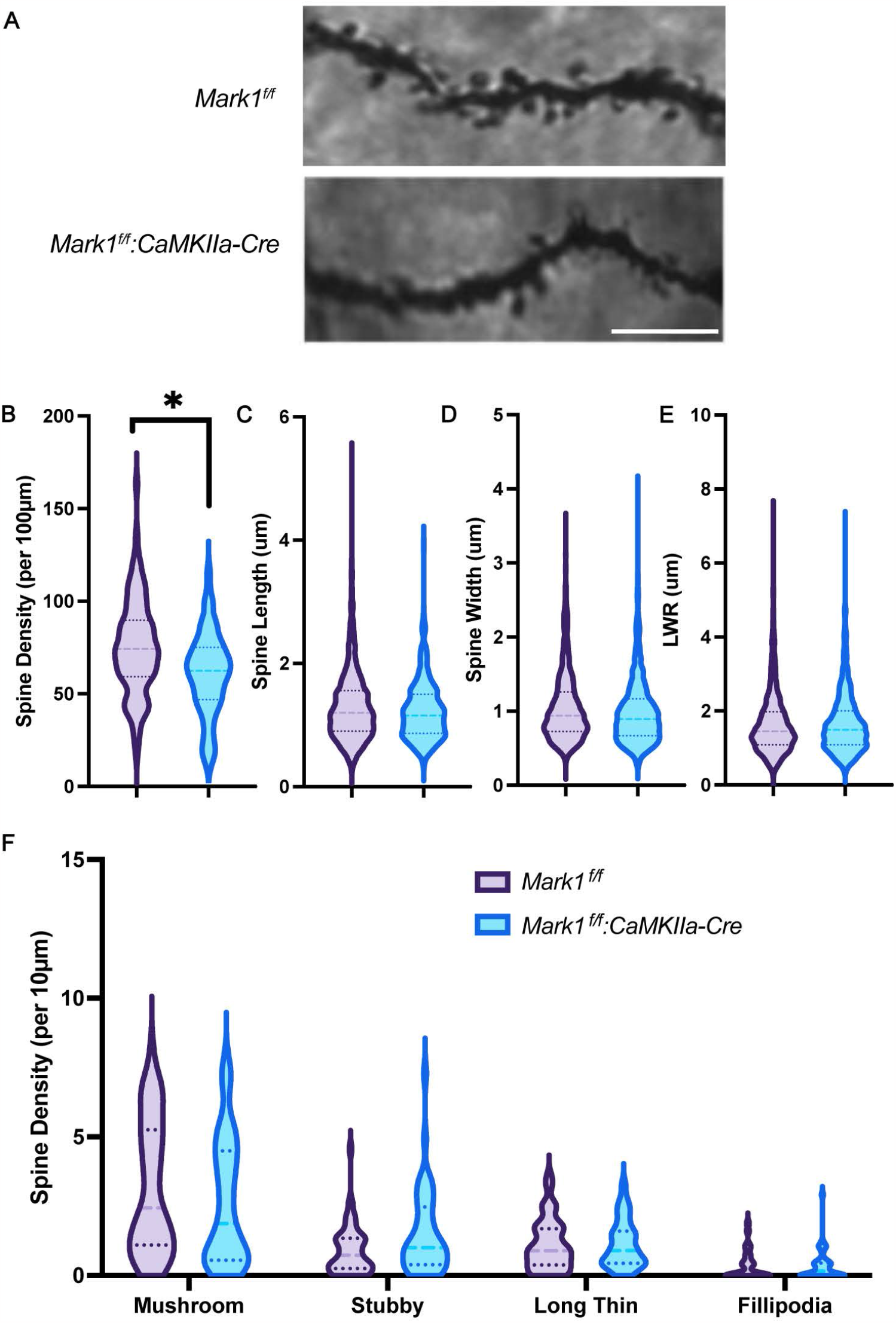
Dendritic spine density is decreased in hippocampal CA1 of MARK1^f/f^:CaMKIIa-Cre mice. A. Image representation of dendritic spines of the *Mark1*^*f/f*^ and *Mark1*^*f/f*^*:CaMKIIa-Cre*. Scale bar: 10μm. B. Graph of mean spine density. n=103 dendrites/5 mice for *Mark1*^*f/f*^, n=89 dendrites/4 mice for *Mark1*^*f/f*^*:CaMKIIa-Cre*, p=0.0172 by mixed model analysis. C. Graph of mean spine length. n=1171 spines/6 mice for *Mark1*^*f/f*^, n=868 spines/4 mice for *Mark1*^*f/f*^*:CaMKIIa-Cre*, p=0.9896 by mixed model analysis. D. Graph of mean spine width. n=1171 spines/6 mice for *Mark1*^*f/f*^, n=868 spines/4 mice for *Mark1*^*f/f*^*:CaMKIIa-Cre*, p=0.4627 by mixed model analysis. E. Graph of mean spine length to width ratio. n=1171 spines/6 mice for *Mark1*^*f/f*^, n=984 spines/4 mice for *Mark1*^*f/f*^*:CaMKIIa-Cre*, p=0.6058 by mixed model analysis. F. Graph of spines classified by morphology. n=1171 spines/59 dendrites/5 mice for *Mark1*^*f/f*^, n=984 spines/46 dendrites/4 mice for *Mark1*^*f/f*^*:CaMKIIa-Cre*. Mushroom p=0.2931, stubby p=0.1115, long thin p>0.9999, filipodia p>0.9999 by 2-way ANOVA.

### Loss of MARK1 leads to an increase in synaptic levels of GKAP and GluR2

Our previous research shows that MARK1 phosphorylates the synaptic scaffolding protein PSD-95 on Ser561. This phosphorylation regulates PSD-95 dynamics and the interaction with its binding partners ^4,27^. To determine whether loss of MARK1 affects the assembly and anchoring of glutamate receptors and other synaptic proteins to the PSD, we examined the levels of glutamate receptors and scaffolding proteins in the total lysate and crude synaptosomal fractions of WT and MARK1 cKO hippocampi. Interestingly, we found that synaptic levels of GluR2 and the PSD-95 interacting partner GKAP are significantly increased in the MARK1 cKO hippocampi as compared with littermate WT controls. No significant changes in the synaptic levels of GluR1, NMDAR1, NMDAR2B, PSD-95, SHANK3, and mGluR5 were observed (**Figure 3A**). A small but statistically significant increase in PSD-95 levels was also observed in the total hippocampal lysate from the MARK1 cKO mice (**Figure 3B**).

**Figure 3:**
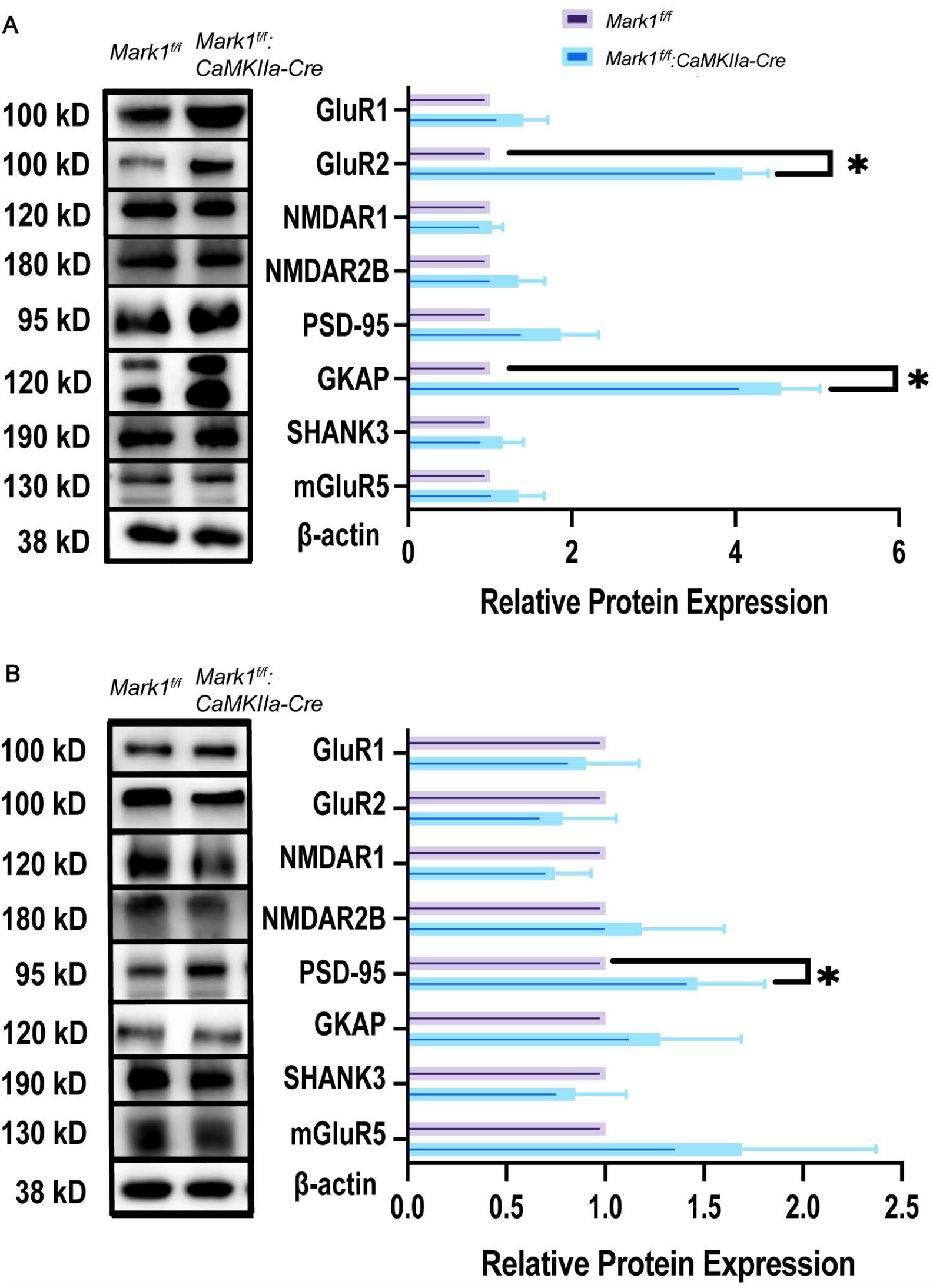
Mark1^f/f^:CaMKIIa-Cre mice display a significant increase of GKAP and GluR2 of crude synaptosome fraction. A. Representative blots and quantification for proteins detected in the hippocampus crude synaptosome fraction from *Mark1*^*f/f*^ and *Mark1*^*f/f*^*:CaMKIIa-Cre* mice. Data is shown as mean ± SEM, n=4-6 mice per protein per genotype. p<0.05 by Unpaired t test with Welch’s correction. B. Representative blots and quatification for proteins detected in the hippocampus homogenate fraction from *Mark1*^*f/f*^ and *Mark1*^*f/f*^*:CaMKIIa-Cre* mice. Data is shown as mean ± SEM, n=4-6 mice per protein per genotype. p<0.05 by Unpaired t test with Welch’s correction.

### Loss of MARK1 impairs spatial learning in the Morris Water Maze

Given the defects in dendritic spine morphogenesis and synaptic transmission observed in the MARK1 cKO mice, we next examined whether hippocampal dependent learning and memory are affected in these mice. The Morris Water Maze task is a test for spatial learning and memory that is highly dependent on the hippocampus, including the CA1 pyramidal neurons. WT and MARK1 cKO mice between 4-5 weeks old were trained in this task and their latency to reach the hidden platform was recorded. We observed that the MARK1 cKO mice have significantly impaired learning in this task compared with littermate controls **(Figure 4B)**. On Day 7, the platform was removed, and the spatial memory was measured by recording the amount of time spent swimming in the target quadrant **(Figure 4C**). No significant differences were observed between WT and MARK1 cKO mice. These results suggest that MARK1 cKO mice do not show any obvious spatial memory deficits even though they show impaired spatial learning.

**Figure 4:**
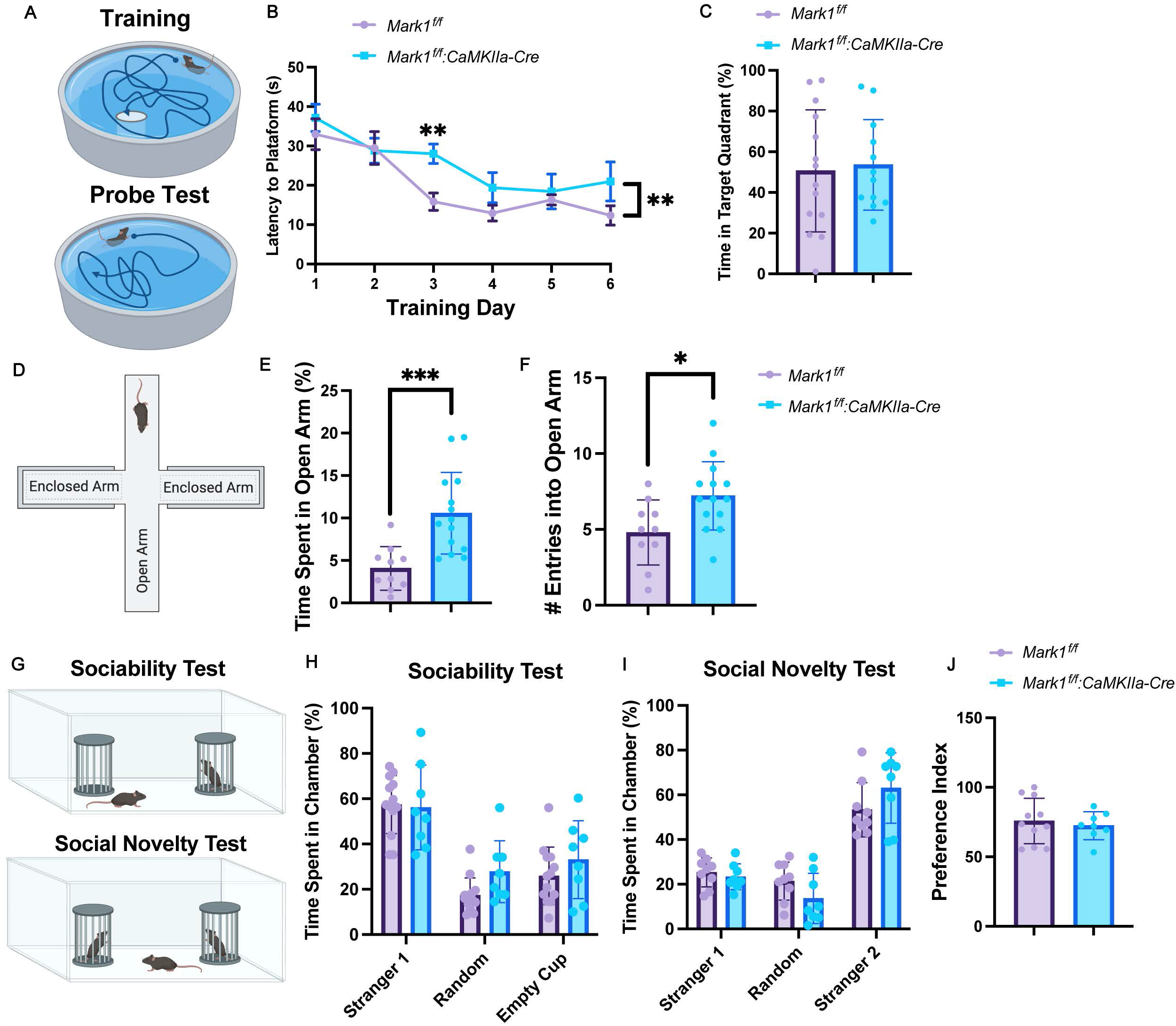
Mark1^f/f^:CaMKIIa-Cre mice show a slower learning curve in the MWM task and reduced anxiety-like behavior in the EPM. A. Graphical representation of the experimental design used for the Morris Water Maze task. B. Graphed average latency to platform(s) of male mice by day. Data shown as mean ± SEM, n=14 for *Mark1*^*f/f*^, n=13 for *Mark1*^*f/f*^*:CaMKIIa-Cre*, p=0.0052 by 2-way ANOVA. C. Graphed average % time spent on targeted quadrant of male mice. Data shown as mean ± SEM, n=14 for *Mark1*^*f/f*^, n=13 for *Mark1*^*f/f*^*:CaMKIIa-Cre*, p>0.05 by Unpaired t test. D. Graphical representation of the experimental design used for the Elevated Plus Maze. E. Graph of mean percentage of time spent in the open arm. Data is shown as mean ± SEM, n=10 *Mark1*^*f/f*^, n=14 *Mark1*^*f/f*^*:CaMKIIa-Cre*, p=0.0004, Unpaired t test with Welch’s correction. F. Graph of mean number of entries into the open arm. Data is shown as mean ± SEM, n=10 *Mark1*^*f/f*^, n=14 *Mark1*^*f/f*^*:CaMKIIa-Cre*, p=0.015, Unpaired t test with Welch’s correction. G. Graphical representation of the experimental design used for the Sociability Test and Social Novelty Test. H. Sociability Test: Mean percentage of time spent in each chamber. Data shown as mean ± SEM, n=13 for *Mark1*^*f/f*^, n=8 for *Mark1*^*f/f*^*:CaMKIIa-Cre*, p>0.05 by Unpaired t test. I. Social Novelty Test: Mean percentage time spent in each chamber. Data shown as mean ± SEM, n=11 for *Mark1*^*f/*f^, n=8 for *Mark1*^*f/f*^*:CaMKIIa-Cre*, p>0.05 by Unpaired t test. J. Mean preference index. Data shown as mean ± SEM, n=11 for *Mark1*^*f/f*^, n=8 for *Mark1*^*f/f*^*:CaMKIIa-Cre*, p>0.05 by Unpaired t test.

### MARK1 cKO mice show reduced anxiety-like behaviors in the Elevated Plus Maze

Since MARK1 is associated with ASD, we next aimed to determine whether loss of forebrain MARK1 leads to behavioral defects relevant to ASD, such as social and anxiety-related behaviors. Thus, we first analyzed anxiety-like behaviors using the Elevated Plus Maze (EPM). This task consists of a plus-shaped maze with two long arms with tall walls and two short arms with no walls (**Figure 4D**). Mice will naturally avoid open and elevated spaces since they might represent a danger to them^34^. The number of times they entered the open arm and the percentage of time spent on the open arm were recorded (**Figure 4E and F**). We found MARK1 cKO mice showed a significant increase in the number of times entering the open arm, as well as the total amount of time spent on the open arm (**Figure 4E and F**). These results show that MARK1 cKO mice exhibit reduced anxiety-like behavior.

### No defects in social behavior in MARK1 cKO mice

Next, we examined social behaviors in the MARK1 cKO mice by using sociability and social novelty tests. Each experimental mouse was exposed to an age- and sex-matched stranger mouse and an empty cup for 10 minutes for the sociability test (**Figure 4G**). We found no significant difference between WT and MARK1 cKO mice in their sociability (**Figure 4H**).

To further investigate the social behavior of our Par1c/MARK1 cKO mice, we conducted the social novelty test 3 hours after the sociability test. To perform this assay, the empty cup from the sociability test was replaced with a second stranger mouse (**Figure 4G**). The amount of time the experimental mouse spent interacting with Stranger 1 (familiar) or Stranger 2 (novel) was recorded (**Figure 4I and J**). Both WT and MARK1 cKO mice spent more time exploring the novel Stranger 2 mouse, and no significant differences were observed between the two genotypes (**Figure 4I and J**). These data suggest that conditionally knocking out MARK1 from forebrain pyramidal neurons does not significantly affect social behaviors in mice.

## Discussion

In this study, we used a novel forebrain-specific MARK1 cKO mouse line to investigate the *in vivo* role of MARK1 in the brain. We found loss of MARK1 in the forebrain pyramidal neurons causes delays in spatial learning in the Morris Water Maze. This is in line with our observation that MARK1 cKO mice show a reduction in dendritic spine density in hippocampal CA1 pyramidal neurons. Interestingly, *MARK1* shows accelerated evolution in the human lineage, and human brains show increased density, which is believed to contribute to our enhanced learning capacity. It will be of great interest to examine how human MARK1 differentially regulates dendritic spines as compared with MARK1 from other mammalian species.

*MARK1* is a Category 2 SFARI gene and is considered a strong candidate for ASD. In addition, MARK1 is also associated with bipolar disorders^30^. Both disorders are often comorbid with anxiety. Thus, it is interesting that we observed a decrease in anxiety-like behavior in the Elevated Plus Maze. However, it is important to note that it remains unclear how MARK1 activity is affected in ASD and bipolar disorders. In fact, one of the ASD-linked SNPs in *MARK1* modulates the transcription of the *MARK1* gene, and *MARK1* was found to be overexpressed in the brains of autistic subjects^28^. Thus, it is possible that MARK1 activation promotes anxiety-like behaviors, and loss of MARK1 would lead to decreased anxiety. Intriguingly, although social dysfunction is classically observed in both ASD and bipolar disorder, we did not observe any defects in social behavior in the forebrain-specific MARK1 cKO mice. MARK1 has widespread high expression throughout many brain regions. Thus, it remains possible that MARK1 may contribute to social behaviors in other brain regions such as the striatum.

We further analyzed synaptic changes in the MARK1 cKO hippocampi and observed a significant increase in synaptic levels of GKAP and GluR2. The increase in synaptic GKAP is in line with our previous studies showing that MARK1 phosphorylation of PSD-95 on Ser561 affects its interaction with GKAP^27^. A non-phosphorylatable mutation of Ser561 promotes PSD-95 interaction with GKAP. Thus, loss of MARK1 would be expected to increase PSD-95 interaction with GKAP, which can potentially lead to an increase in synaptic levels of GKAP.

The increase in synaptic GluR2 is unexpected, especially since this increase seems specific to GluR2, with no accompanying change in GluR1. It is possible that the increased synaptic GKAP helps stabilize GluR2 at the spine surface, although further studies are needed to examine the underlying mechanism by which GluR2 is specifically stabilized at the synapses. Alternatively, MARK1 may affect a yet unknown target to increase GluR2 trafficking to the spine surface. Overall, these findings suggest that the loss of MARK1 leads to alterations in synaptic protein composition and organization, potentially contributing to the observed defects in dendritic spine morphogenesis and cognitive functions.

Taken together, our studies show a novel role of MARK1 in regulating dendritic spine morphogenesis and cognitive functions *in vivo*. Further investigations are warranted to elucidate the precise molecular mechanisms by which MARK1 regulates synaptic plasticity and cognitive functions, as well as its potential implications in the pathogenesis of neurodevelopmental and psychiatric disorders.

## Acknowledgements

This work was supported by NIH grants NS089578 and NS134233 to HZ. EK was supported by the Rutgers IMSD Pipeline program (NIH R25GM055145) and a diversity supplement to NS089578. We would like to thank Dr. George Wagner for his expert advice on the behavioral studies.

## Notes

### Competing Interest Statement

The authors have declared no competing interest.

