## Supplemental Information for "MARK1 regulates dendritic spine morphogenesis and cognitive functions *in vivo*"

***Supplemental Figure 1: MARK1 conditional knockout mouse model shows no differences in body or brain weight and no differences in MARK2 or MARK3 expression.***

A. Graph of mean body weight (g). Data is shown as mean ± SEM, n=12 for *Mark1$^{f/f}$* (n=8 males, n=4 females), n=13 for *Mark1$^{f/f}$:CaMKIIa-Cre* (n=6 males, n=7 females), p>0.05 by Unpaired t test.
B. Graph mean whole-brain weight (g). Data is shown as mean ± SEM, n=9 for *Mark1$^{f/f}$* (n=6 males, n=3 females), n=7 for *Mark1$^{f/f}$:CaMKIIa-Cre* (n=3 males, n=4 females), p>0.05 by Unpaired t test.
C. Representative blots and quatification for MARK2 and MARK3 proteins detected in the hippocampus crude synaptosome fraction from *Mark1$^{f/f}$* and *Mark1$^{f/f}$:CaMKIIa-Cre* mice. Data is shown as mean ± SEM, n=4-6 mice per protein per genotype.

Supplemental Figure 1

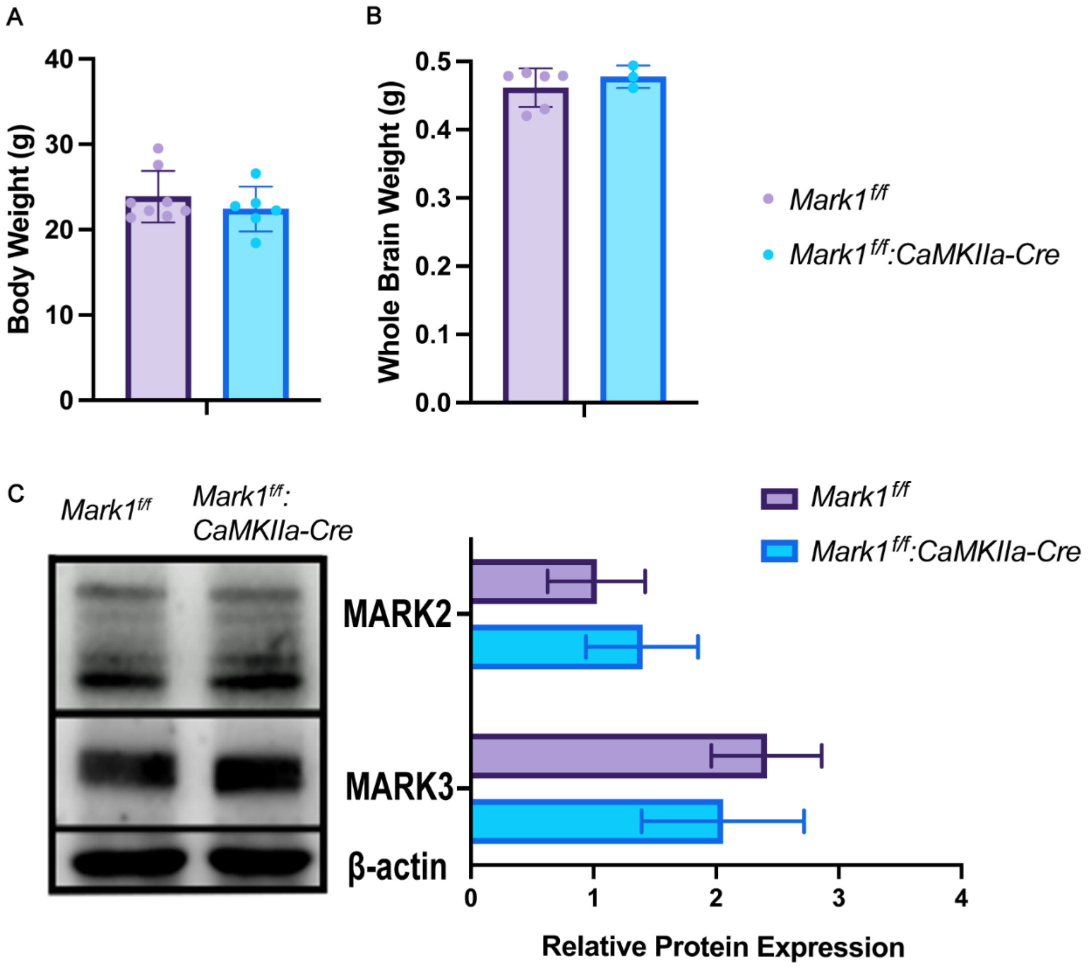
